# [^18^F]Flortaucipir binding and affinity to tau protein measured in postmortem human brains of Alzheimer’s disease and Down syndrome by autoradiography: Comparison with [^18^F]MK-6240

**DOI:** 10.64898/2026.11.15.766816

**Authors:** Agnes P. Biju, Josie M. Herrera, Chloe C. McNamara, Emma Lynn M. Hipolito, Christopher Liang, Jogeshwar Mukherjee

**Author notes:** (A.P.B); (J.M.H); (C.C.M); (E.M.H); (C.L.); (J.M.). Address Correspondence to: Jogeshwar Mukherjee, Ph.D. Medical Sciences B138, Department of Radiological Sciences, University of California, Irvine, Irvine, CA 92697-5000.

## Abstract

Flortaucipir and MK-6240 are two tau imaging agents currently in use for measuring neurofibrillary tangle (NFT) progression in Alzheimer’s disease (AD). The aims of this study were to use postmortem AD and Down syndrome (DS) brain sections to study three issues: (1) Measure binding affinity of flortaucipir and MK-6240 in AD brain sections by using [^125^I]IPPI; (2) Evaluate the binding characteristics of [^18^F]flortaucipir in AD, DS and cognitively normal (CN) brains sections and compare them with [^18^F]MK-6240, and (3) Assess the binding of [^18^F]flortaucipir in CN and Parkinson’s disease (PD) brain sections to identify any off-target binding. The half-maximal inhibition (IC50) of [^125^I]IPPI bound to anterior cingulate (AC) in AD brain sections for flortaucipir was 2.91 nM and MK-6240 was 5.99 nM. Using frontal cortex (FCX) and temporal cortex (TCX) of AD, DS and CN brain sections, [^18^F]flortaucipir displayed high levels of binding in the gray matter (GM) of AD and DS. The average ratio of DSAD GM to CN GM was 15.7 in FCX, while DSAD GM to CN GM was 15.2 for the TCX. In the case of AD, average ratio of AD GM to CN GM was 12.4 in FCX, while AD GM to CN GM was 9.5 for the TCX. Binding of [^18^F]flortaucipir in brain section of AC from CN and PD cases did not exhibit any off-target binding to monoamine oxidase-A (MAO-A). In summary, both flortaucipir and MK-6240 exhibit similar high affinity to tau and [^18^F]flortaucipir exhibits higher binding compared to [^18^F]MK-6240 in DSAD.

## 1. Introduction

Down syndrome (DS) associated Alzheimer’s disease (AD) is a progressive neurodegenerative disorder pathologically attributed to the accumulation of fibrillar amyloid beta (Aβ) plaque as well as tau neurofibrillary tangles (NFT) (<u>Fortea et al., 2021</u>, <u>Zammit et al., 2024</u>; <u>Rafii et al., 2025</u>). Trisomy 21, the triplication of APP gene on chromosome 21 resulting in the Aβ plaque and NFT, makes DS genetically similar to AD (<u>Fortea et</u> <u>al., 2021</u>; <u>Wisch et al., 2025</u>). Extensive studies in AD suggest that NFT contribute largely to cognitive decline, more so than amyloid plaques, leading them to be an imperative marker in predicting disease progression (<u>Huber et al., 2017</u>; <u>Malpetti et al., 2022</u>).

Two NFT-Positron emission tomography (PET) imaging agents have now been widely used in AD. [^18^F]Flortaucipir (also referred to as T807 or AV-1451) and [^18^F]MK-6240, two PET radiotracers, belonging to different classes of compounds, are being widely used to detect NFT (<u>Lowe et al., 2016</u>; <u>Li et al., 2021</u>, <u>Malarte et al., 2021</u>; <u>Mukherjee et al., 2021</u>). Both PET radiotracers have successfully mapped tau burden in high-vulnerability brain regions in AD subjects with some off-target issues of both the radiotracers (<u>Choi et</u> <u>al., 2018</u>; <u>Gogola et al., 2022</u>; <u>Tissot et al., 2023</u>; <u>Flores et al., 2025</u>; <u>McVea et al., 2024</u>). Cognitive decline throughout the progression of the disease is associated with changes in NFT supported by tau PET radiotracers depicting tau pathology in these regions (<u>Bischoff et al., 2023</u>). Comparative studies of [^18^F]flortaucipir and [^18^F]MK-6240 continue to find subtle differences in the two radiotracers related to early stage and late stage tau pathology detection (<u>Povola et al., 2026</u>).

Postmortem studies comparing DSAD and AD show very similar levels of Aβ plaque and NFT (<u>Lemoine</u> <u>et al., 2020</u>; <u>Biju et al., 2026</u>). Both [^18^F]flortaucipir and [^18^F]MK-6240 have now advanced to PET studies of DSAD subjects (<u>Zammit et al., 2024</u>; <u>Zammit et al., 2026</u>) and increased tau has been related to lower cognition in DSAD (<u>Zammit et al., 2026</u>). Efforts have been made in postmortem DSAD and AD brains to understand off-target binding of these tracers (<u>Karim et al., 2026</u>; <u>Aguero et al., 2024</u>). The sporadic meningeal uptake of [^18^F]MK-6240 has been observed in both AD and DSAD subjects (<u>McVea et al., 2024</u>; <u>Zammit et al., 2026</u>) and efforts have been made to characterize this non-tau binding (<u>Karim et al., 2026</u>). Several studies on [^18^F]flortaucipir have been reported in an effort to understand off-target retention in the choroid plexus and basal ganglia (<u>Choi et al., 2018</u>) and potential monoamine oxidase (MAO) binding (<u>Vermeiren et al., 2018</u>; <u>Drake et al., 2019</u>; <u>Hansen et al., 2018</u>; <u>Wright et al., 2022</u>). Structurally, [^18^F]flortaucipir has some resemblance to harmine, a predominantly MAO-A binding drug with some tau affinity (McNamara et al., 2026; <u>Karim et al., 2025</u>). Although high nanomolar affinity of flortaucipir to MAO-A (expressed in human recombinant cell lines) has been reported (<u>Vermeiren et al., 2018</u>), identification of its MAO binding in vivo has remained elusive. Thus, it remains unclear if the small differences seen in PET studies of [^18^F]flortaucipir and [^18^F]MK-6240 comes from off-target or pharmacological characteristics of these two commonly used tau radiotracers (<u>Gogola et al., 2022</u>; <u>Povola et al., 2026</u>).

The three objectives of this study were as follows: (1) Measure and compare the binding affinity of MK-6240 and flortaucipir autoradiographically in AD brain slices by using the same selective tau radioligand, [^125^I]IPPI (<u>Mukherjee et al., 2021</u>; <u>Karim et al., 2025</u>); (2) Compare the binding properties of [^18^F]flortaucipir with our recent findings of [^18^F]MK-6240 in DSAD, AD, and cognitively normal (CN) cohorts (<u>Karim et al.,</u> <u>2026</u>). Frontal cortex (FCX) and temporal cortex (TCX) from the cases were used for the study, and (3) Conduct preliminary autoradiographic study of [^18^F]flortaucipir in Parkinson’s disease (PD) anterior cingulate brain slices. We have previously reported high levels of MAO-A binding using [^18^F]FAZIN3 in PD (<u>Mukherjee</u> <u>et al., 2022</u>) and AD (<u>Syed et al., 2023</u>). The use of PD brain tissue sections minimizes potential interference due to the absence of [^18^F]flortaucipir tau binding (<u>Marquie et al., 2017</u>; <u>Burnham et al., 2024</u>).

## 2. Materials and Methods

### 2.1. Human Tissue Samples

Institutional Biosafety Committee of University of California, Irvine approved acquisition and studies using post-mortem human brain tissue. Postmortem brain tissue samples of DSAD (n=5), AD (n=5), and CN (n=5) cases (male and female), each consisting of FCX and TCX, were obtained from UCI Memory Impairment and Neurological Disorders (MIND) institute for in vitro experiments (Table 1). Anterior cingulate brain tissue used in this study were obtained from Banner Health Research Institute (BHRI), Sun City, Arizona (AD=3; CN=2; PD=6, Table-1; <u>Beach et al., 2015</u>). Brain slices, 10 µm thick, were obtained from the frozen tissue blocks using a Leica 1850 cryotome cooled to –20 °C and collected on Fisher glass slides. All slides were then stored at –80 °C. Neighboring brain slices (within 200 microns) from the tissue block of each case were used for various in vitro studies.

**Table 1.**
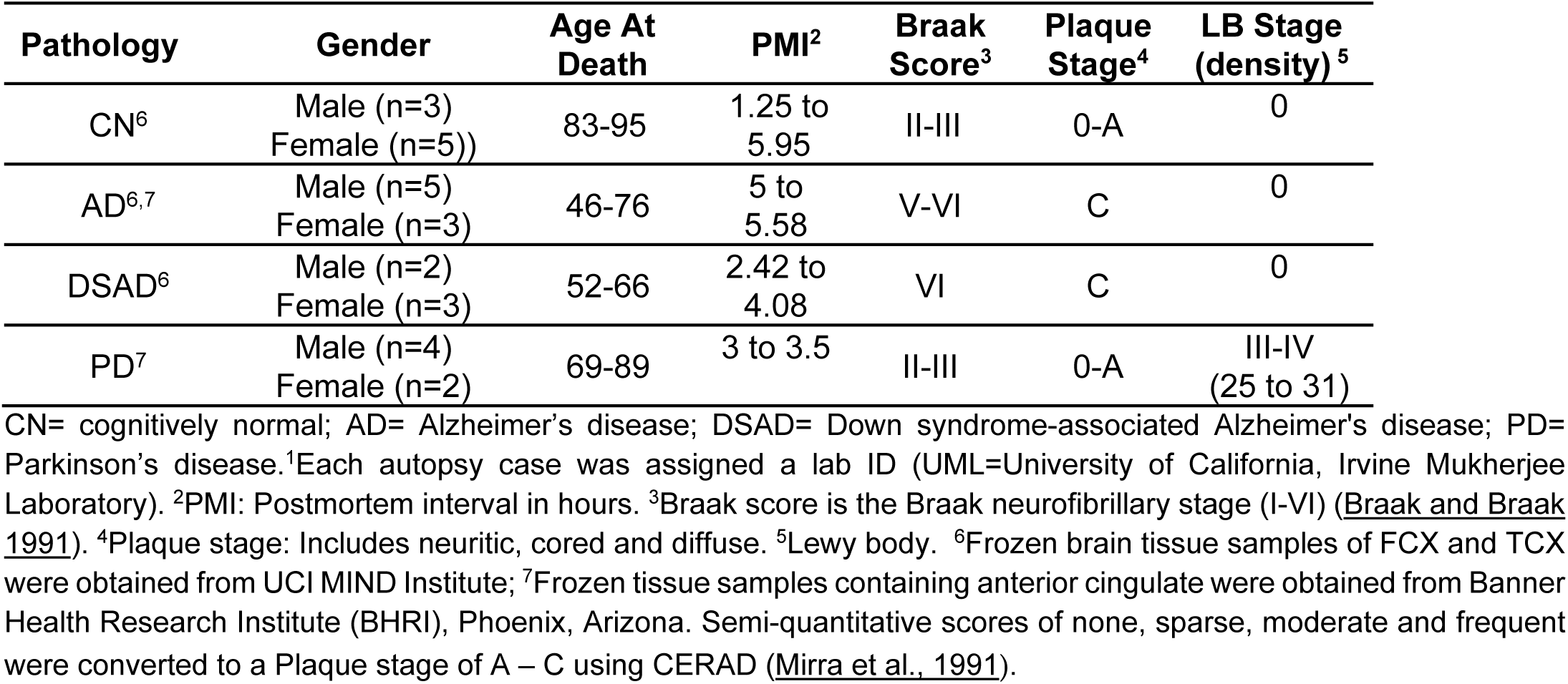
Brain tissue cases and data.

Brain slices of all cases were immunostained for tau by UCI Pathology core services as previously described (<u>Karim et al., 2026</u>; <u>Biju et al., 2026</u>). For total Tau (<u>Sjogren et al., 2001</u>), DAKO polyclonal antibody, which detects all 6 six isoforms of tau from the microtubule-associated protein tau gene, was used at a dilution 1:3000, A0024 (Agilent, CA, USA). Immunostained slides were scanned for analysis using QuPath (v 0.7.0) protocol (<u>Mondal et al., 2023)</u>.

### 2.2. Radiopharmaceuticals

Flortaucipir (also referred as AV-1451 or T807 in the manuscript) and the precursor for [^18^F]flortaucipir, *tert*-butyl-7-(6-nitropyridin-3-yl)-5H-pyrido[4,3-b]indole-5-carboxylate were purchased from AA Blocks, Inc., San Diego, USA. Using high specific activity fluorine-18 from PETNET, Culver city, CA, [^18^F]flortaucipir (also referred as [^18^F]AV-1451 or [^18^F]T807 in the manuscript) was prepared using reported methods with >95% radiochemical purity and specific activity of 74 GBq/μmol (<u>Mossine et al., 2017</u>). Specific activity of [^18^F]flortaucipir was comparable to the specific activity of [^18^F]MK-6240 used in our recent study (<u>Karim et al.,</u> <u>2026</u>). ^125^I]IPPI was prepared using our previously reported methods of radioiodinations of this class of compounds (<u>Mukherjee et al., 2021</u>; <u>Limpengco et al., 2023</u>; <u>Sison et al., 2024</u>). Unlabeled MK-6240 was purchased from1ClickChemistry, Inc, Tinton Falls, New Jersey, USA. All other biochemicals, reagents and solvents were purchased from Sigma-Aldrich, Inc, St. Louis. Flortaucipir and MK-6240 were dissolved in absolute ethanol and used for competition studies.

### 2.3. [^125^I]IPPI Autoradiography for Binding Affinity

Binding affinity measurements of flortaucipir and MK-6240 were carried out by using [^125^I]IPPI autoradiography. Brain slices (10 μm thick) containing anterior cingulate (AC) from 3 AD cases were used to evaluate change in [^125^I]IPPI specific binding at a concentration range of 10^-9^, 10^-8^, 10^-7^, 10^-6^ and 10^-5^ M of flortaucipir and MK-6240. Slides were placed in Fisher brand chambers (volume 20 mL) and preincubated with phosphate buffered saline (PBS) buffer at ambient temperature for 15 minutes. The preincubation buffer was discarded and [^125^I]IPPI (3.7 kBq/mL) in PBS buffer was added to all the chambers (total binding, nonspecific binding with 10^-4^M MK-6240 and chambers containing the two drugs at different concentrations) and incubated for one hour. After incubation, the slides were washed with 100% PBS (1 minute), 70% ethanol in PBS (2 minutes), 30% ethanol in PBS (1 minute), and lastly 100% PBS for 1 minute. Washed slides were air-dried and apposed to phosphor films in cassettes for 5 to 7 days. The phosphor films were read on the Cyclone Phosphor Imager, and the images were analyzed using the acquisition and analysis OptiQuant software (Packard Instruments Co, version 5.0). Region-of-interest (ROI) included the gray matter region of AC, which contained NFT and white matter region of corpus callosum which lacked NFT and binding was measured in digital light unit per unit mm^2^ (DLU/mm^2^). Nonspecific binding (10^-4^M MK-6240) was subtracted from total binding from all conditions to provide specific binding. GraphPad Prism (v 11.1.0) was used to compute half-maximal inhibitor concentration (IC_50_) of flortaucipir and MK-6240.

### 2.4. [^18^F]Flortaucipir Autoradiography

Brain sections of all DSAD, AD and CN cases included FCX and TCX on separate slides with up to 3 brains sections on each slide. Preincubation of all the slides was carried out for 15 minutes in PBS buffer (6 slides per glass chamber, total volume 60 mL). A stock PBS pH 7.4 buffer solution of [^18^F]flortaucipir (∼148 kBq/mL) was used in all the chambers. This concentration of [^18^F]flortaucipir was lower than [^18^F]MK-6240 used in our recently reported study (<u>Karim et al., 2026</u>). The chambers were incubated at ambient temperature for 1 hour and then underwent multiple washes as described above for [^125^I]IPPI, which was similar to the [^18^F]MK-6240 protocol (<u>Karim et al., 2026</u>). The brain sections were air dried and then transferred into a film cassette with a phosphor screen film. After 24 hours, the films were read on the Cyclone Phosphor imager (Packard Instruments Co). OptiQuant was used to draw regions of interest ROIs on the autoradiographic images, and the extent of binding was measured in DLU/mm^2^. Background activity levels were subtracted from all images. Region-of-interest (ROI) included the gray matter (GM) regions of FCX and TCX which contained NFT and white matter (WM) regions which lacked NFT and binding was measured in digital light unit per unit mm^2^ (DLU/mm^2^). These regions were confirmed by the anti-tau immunostained sections of each case. Higher DLU/mm^2^ from autoradiography indicated higher [^18^F]flortaucipir binding.

[^18^F]Flortaucipir binding in AC brain sections from CN and PD brain cases was similarly carried out as described above. Region-of-interest (ROI) included the GM region of AC and WM region of corpus callosum. Anti-ubiquitin and anti-α-synuclein immunostained sections of each case were used for ROI delineation as described previously for the [^18^F]FAZIN3 study in the same cases (<u>Mukherjee et al., 2022</u>). Clorgyline (AA Blocks, Inc., San Diego, USA), an irreversible MAO-A binding drug was used for competition experiments with [^18^F]flortaucipir, similar to the reported experiments with [^18^F]FAZIN3 study in the same CN and PD cases (<u>Mukherjee et al., 2022</u>).

### 2.5. Statistical Analysis

[^18^F]Flortaucipir and [^125^I]IPPI DLU/mm^2^ values from OptiQuant were analyzed in GraphPad Prism 11.1 (GraphPrism Software, CA, USA) and Microsoft Excel 16. Student’s t-test were performed with p values <0.05 indicating statistical significance. Error bars are presented as mean ± standard deviation. Pearson’s correlation and linear regression was used for the parametric comparison between [^18^F]flortaucipir DLU/mm^2^ values within DSAD and AD cases were determined in GraphPad Prism 11.1.

## 3. Results

### 3.1 In Vitro Binding Affinity

Using the same radioligand, [^125^I]IPPI, we measured the binding affinity of the two structurally different compounds, flortaucipir (Fig-1A) and MK-6240 (Fig-1B). The 3 AD cases (BML72, BML62 and BML89) chosen for the autoradiographic study were all Braak V stage and had extensive amount of tau as shown in the immunostains (Fig-1C,D,E). As expected, very selective binding of [^125^I]IPPI was observed in the AC brain sections of all the 3 AD subjects as seen in Fig. 1, similar to our previous reports (<u>Mukherjee et al., 2021</u>; Mondal et al., 2023). MK-6240 (100 μM) was able to displace >98% of [^125^I]IPPI from all the brain slices and was used as a measure of nonspecific binding (Fig-1I,J,K). [^125^I]IPPI binds to human tau with an inhibition constant, Ki=0.75 nM, and belongs to the same class of compounds, azaindoles, with some structural resemblance to MK-6240 (<u>Mukherjee et al., 2021</u>). Increasing the concentration (from 10^-9^ to 10^-5^M) of flortaucipir and MK-6240 progressively reduced the specific binding (total-nonspecific) of [^125^I]IPPI to the AC of the 3 AD cases as seen in Fig-1L,M.

**Fig. 1.**
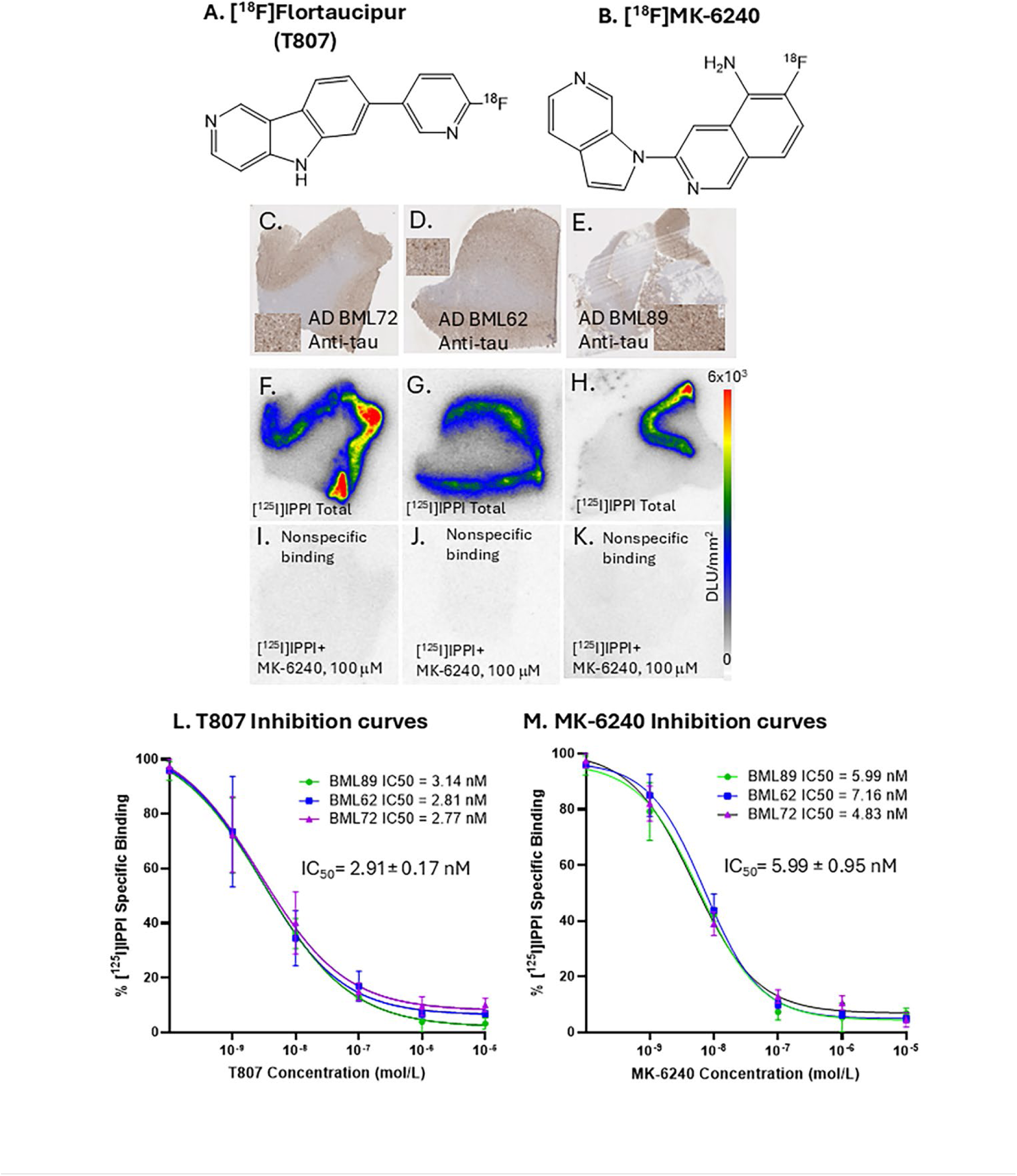
In vitro binding affinity to tau in AD: (A). Chemical structure of [^18^F]flortaucipir; (B). Chemical structure of [^18^F]MK-6240; (C,D,E). Anti-tau IHC of postmortem human brain slice (10 μm) of AD BML72, AD BML62 and AD BML89, with inset of 50 µm magnification in each; (F,G,H). [^125^II]IPPI binding to AD BML72, AD BML62 and AD BML89 gray matter regions of anterior cingulate; autoradiography scale bar: 0-1500 DLU/mm^2^; (I,J,K). Nonspecific binding in the presence of 100 μM MK-6240 in AD BML72, AD BML62 and AD BML89; (L). Flortaucipir-induced inhibition curves of specific binding of [^125^I]IPPI, average IC_50_=2.91±0.17 nM; (M). MK-6240-induced inhibition curves of specific binding of [^125^I]IPPI, average IC_50_=5.99±0.95 nM.

Measured half-maximal inhibition concentrations, IC_50_ values for flortaucipir in the 3 AD cases were:3.14 nM, 2.81 nM and 2.77 nM, with an average IC_50_= 2.92±0.17 nM. For MK-6240, the IC_50_ values were 5.99 nM, 7.16 nM and 4.83 nM, with an average IC_50_= 5.99±0.95 nM. These results suggest that flortaucipir has approximately two times better affinity for binding to AD tau in brain sections compared to MK-6240.

### 3.2 [^18^F]Flortaucipir Binding in DSAD, AD, and CN Cases

#### 3.2.1. DSAD Cases

All DSAD cases were Braak VI stage and were expected to show high tau binding, confirmed by anti-tau immunostains (Fig-2A,C,E,G). Cortical binding of [^18^F]flortaucipir was seen in FCX and TCX of the DSAD subjects. Figure-2 shows binding of [^18^F]flortaucipir in two DSAD cases (UML11 and UML12). Distinct binding was observed in the GM rich in tau while WM exhibited approximately 14 to 16% of GM as nonspecific binding (Fig-2I). Although FCX exhibited greater average binding of [^18^F]flortaucipir compared to TCX, it was not significant. Ratio of [^18^F]flortaucipir GM/WM was >7.25 for FCX and >5 for TCX. Binding of [^18^F]flortaucipir was higher in all the DSAD cases compared to our previous report of [^18^F]MK-6240 in these cases (<u>Karim et</u> <u>al., 2026</u>). For [^18^F]MK-6240 the average GM/WM ratio in FCX was 3.1 while TCX had a GM/WM ratio of 2.5 in DSAD. Figure-2J shows the GM/WM ratios of [^18^F]flortaucipir and [^18^F]MK-6240 in all the DSAD cases suggesting the similarity of binding of the two radiotracers. [^18^F]Flortaucipir exhibited approximately two times higher GM/WM ratios compared to [^18^F]MK-6240 in both the brain regions.

**Fig. 2.**
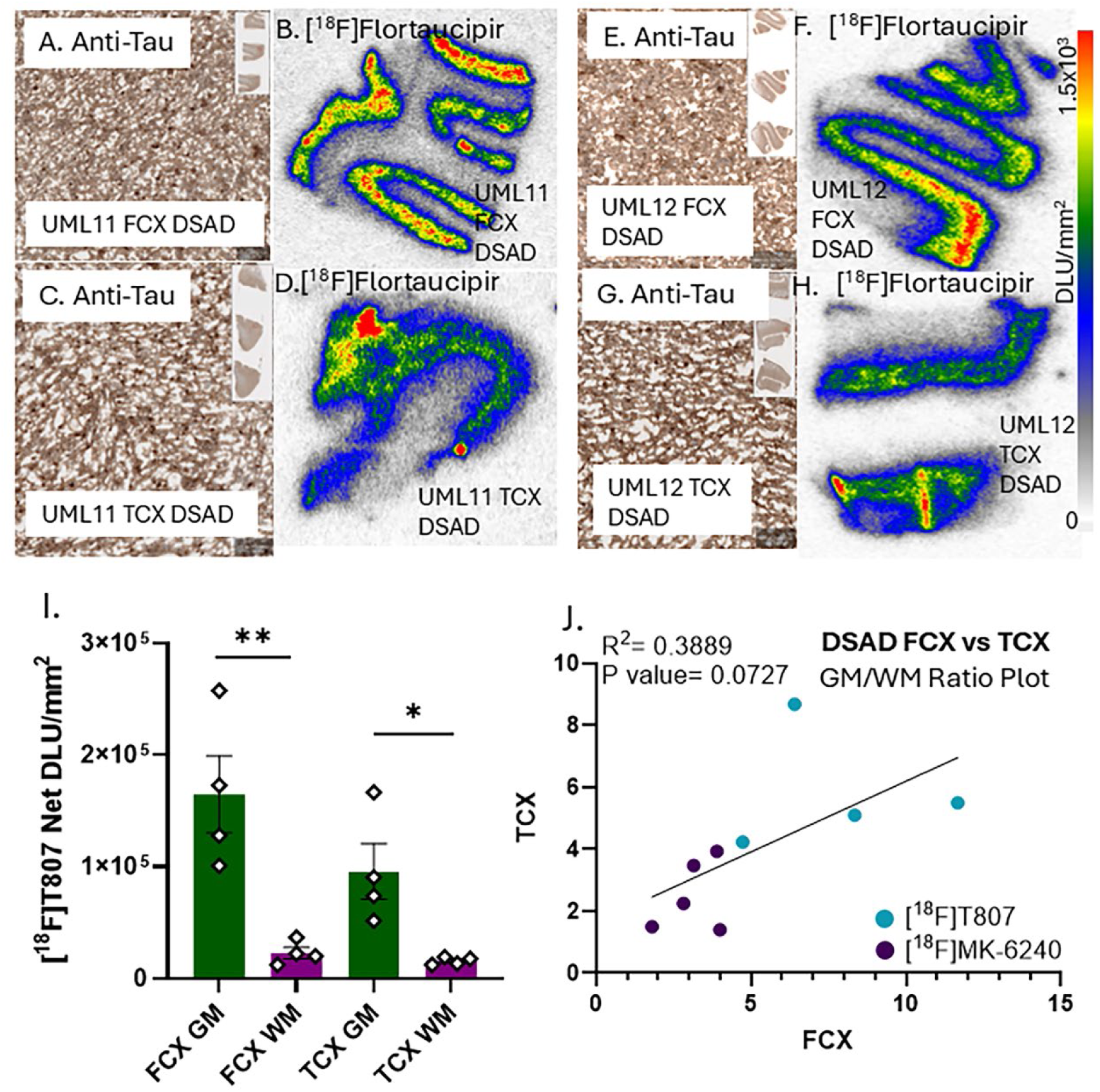
[^18^F]Flortaucipir binding to Tau in DSAD: (A, C, E, G). Postmortem human brain slice (10 μm) of DSAD FCX and TCX of UML11 and UML12 anti-tau IHC of DSAD at 50 µm magnification. (B, D, F. H). [^18^F]Flortaucipir binding to DSAD FCX and TCX of UML11 and UML12, respectively (autoradiography scale bar: 0-1500 DLU/mm^2^). (I). Plot shows binding of [^18^F]flortaucipir in the GM and WM in FCX and TCX of all cases (* p < 0.05, ** p < 0.01). (J). Plot of FCX versus TCX GM/WM ratios for [^18^F]T807 and [^18^F]MK-6240 in the same DSAD cases.

#### 3.2.2. AD Cases

High tau binding was expected based on the anti-tau immunostains in all the AD cases which were of Braak VI stage (Fig-3A,C,E,G). Binding of [^18^F]flortaucipir was seen in FCX and TCX of the AD subjects.

**Fig. 3.**
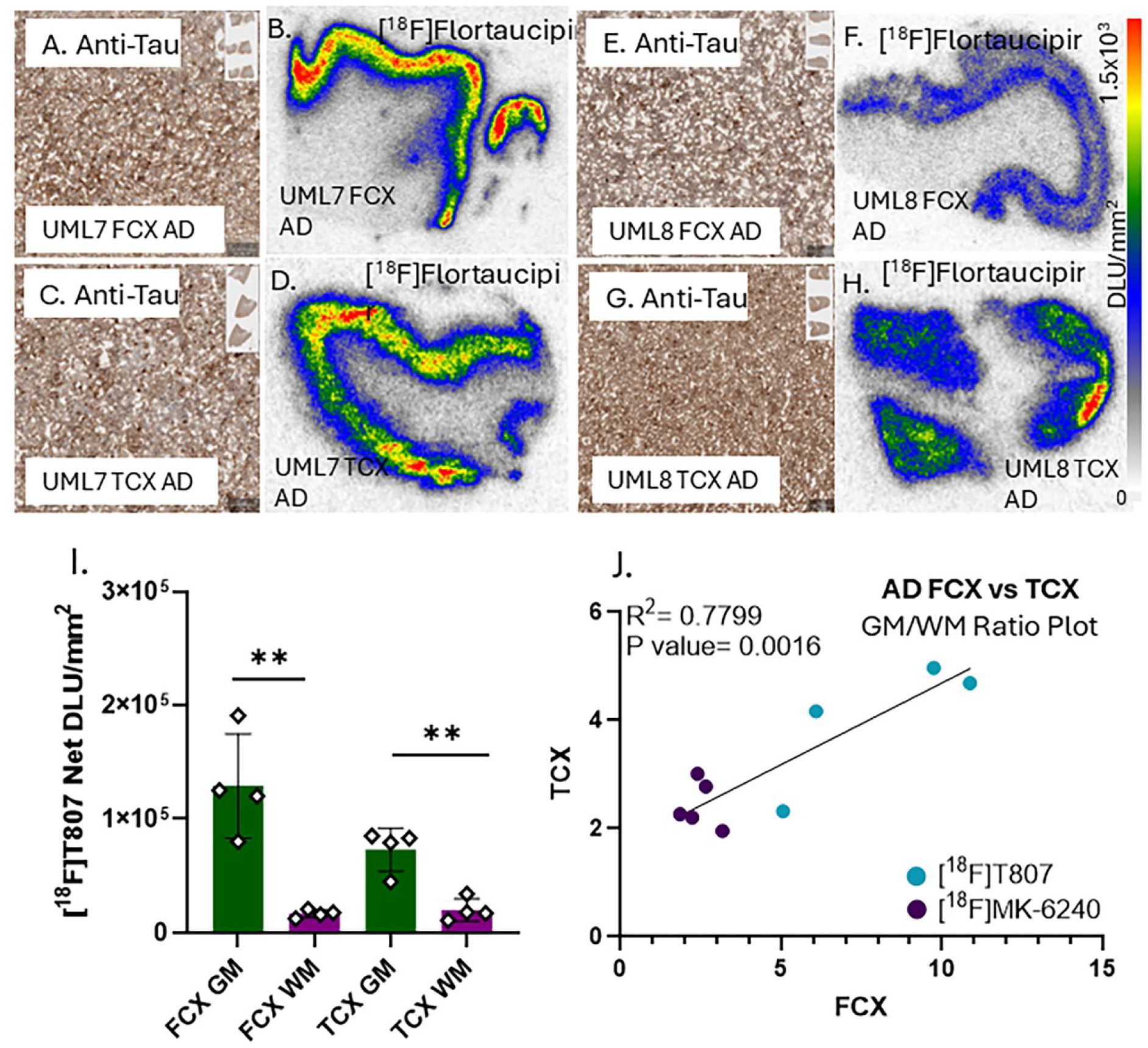
[^18^F]Flortaucipir binding to tau in AD: (A, C, E, G). Postmortem human brain slice (10 μm) of AD FCX and TCX of UML7 and UML8 anti-tau IHC of AD at 50 µm magnification. (B, D, F. H). [^18^F]Flortaucipir binding to AD FCX and TCX of UML7 and UML8, respectively (autoradiography scale bar: 0-1500 DLU/mm^2^). (I). Plot shows binding of [^18^F]flortaucipir in the GM and WM in FCX and TCX GM/WM ratios of all cases. (** p < 0.01). (J). Plot of FCX versus TCX for [^18^F]T807 and [^18^F]MK-6240 in the same AD cases.

Figure-3 shows binding of [^18^F]flortaucipir in two AD cases (UML7 and UML8). Distinct binding was observed in the GM rich in tau was seen while WM exhibited nonspecific binding (Fig-3). Although FCX exhibited greater average binding of [^18^F]flortaucipir compared to TCX but was not significant. Figure-3 shows binding of [^18^F]flortaucipir in two DSAD cases (UML7 and UML8). Distinct binding was observed in the GM rich in tau while WM exhibited ∼3 to 7% of GM as nonspecific binding (Fig-3I). Binding in the FCX of [^18^F]flortaucipir was higher compared to TCX. Ratio of [^18^F]flortaucipir GM/WM was >7.25 for FCX and >4 for TCX in the AD cases. Binding of [^18^F]flortaucipir was higher in all the AD cases compared to our previous report of [^18^F]MK-6240 in these cases (<u>Karim et al., 2026</u>). For [^18^F]MK-6240 the average GM/WM ratio in FCX and TCX had a GM/WM ratio of 2.4-2.5 in AD cases. Thus, [^18^F]flortaucipir exhibited higher GM/WM ratios compared to [^18^F]MK-6240 in both the AD brain regions.

#### 3.2.3. CN Cases

The CN cases were Braak stage II (with two cases in stage III) and were confirmed by the absence of tau in the immunostains (Fig-4A,C,E,G). As expected, [^18^F]flortaucipir did not exhibit any binding to the FCX and TCX of the CN cases (Fig-4B,D,F,H). Plot shows very low levels of [^18^F]flortaucipir in all cases with little difference between GM and WM. The nonspecific binding in the WM of the CN cases were significantly lower (∼25%) compared to the WM nonspecific binding seen in DSAD (Fig-2I) and AD cases (Fig-3I). Absence of significant cortical binding in the CN cases may also suggest the lack of any off-target binding by [^18^F]flortaucipir in these brain regions. Similar observations of a lack of significant cortical binding in the same CN cases were made with [^18^F]MK-6240 (<u>Karim et al., 2026</u>).

**Fig. 4.**
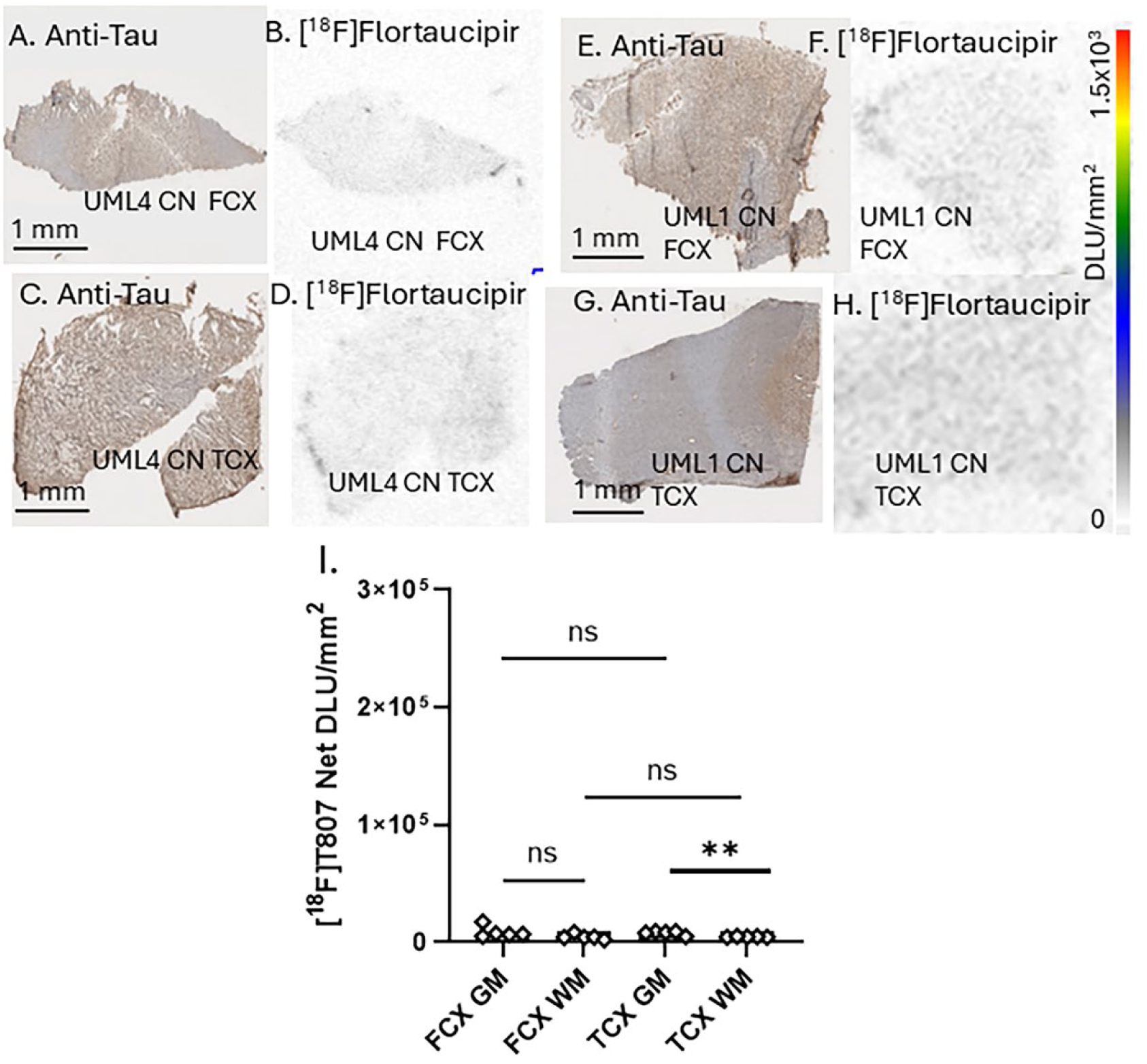
[^18^F]Flortaucipir binding to tau in CN: (A, C, E, G). Postmortem human brain slice (10 μm) of CN FCX and TCX of UML4 and UML1 anti-tau IHC of CN. (B, D, F. H). [^18^F]Flortaucipir binding to CN FCX and TCX of UML4 and UML1, respectively (autoradiography scale bar: 0-1500 DLU/mm^2^). (I). Plot shows binding of [^18^F]flortaucipir in the GM and WM in FCX and TCX of all cases. (** p < 0.01; ns= not significant).

#### 3.2.4. Comparison of DSAD, AD and CN Cases

Comparisons of [^18^F]flortaucipir binding between DSAD, AD, and CN in the GM and WM regions of FCX and TCX are shown in Fig-5. Highest levels of [^18^F]flortaucipir binding were measured in the DSAD cases in the GM regions as seen in FCX (Fig. 5A) and TCX (Fig. 5B). The ratio of DSAD GM to CN GM was 15.7 in FCX, while DSAD GM to CN GM was 15.2 for the TCX. In the case of AD cases, ratio of AD GM to CN GM was 12.4 in FCX, while AD GM to CN GM was 9.5 for the TCX. The average WM in DSAD was the highest (x4.2 in FCX and x2.8 in TCX higher than CN) followed by AD (x3.8 in FCX to x1.3 in TCX higher than CN). This higher WM may be due to spillover effects from GM or there may be traces of tau within these regions.

**Fig. 5.**
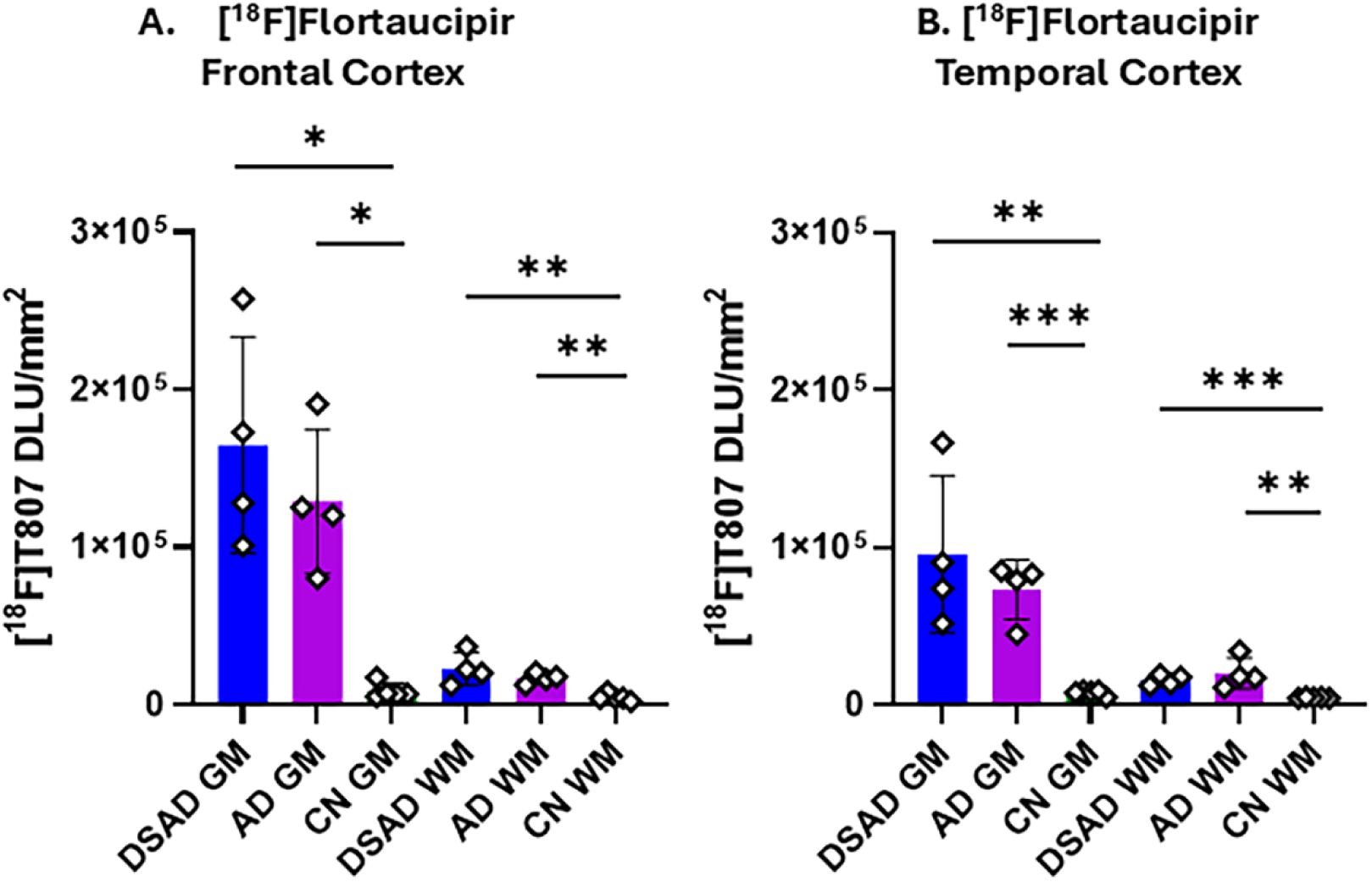
Group Comparisons of [^18^F]Flortaucipir Binding: (A). [^18^F]Flortaucipir binding in FCX GM and WM of DSAD, AD, and CN cases. (B). [^18^F] Flortaucipir binding in TCX GM and WM of DSAD, AD, and CN cases. Unpaired two-tailed parametric t-tests determined statistical significance between each parameter (* p < 0.05, ** p < 0.01, *** < 0.001).

#### 3.2.5. Comparison of [^18^F]flortaucipir and [^18^F]MK-6240

Figure-6A compares the [^18^F]flortaucipir GM/WM ratios of all the FCX and TCX cases in the three groups. The average [^18^F]flortaucipir GM/WM ratios were: DSAD= 6.82, AD= 5.99 and CN=1.70. Figure-6B compares the [^18^F]MK-6240 GM/WM ratios of all the FCX and TCX cases in the three groups reported in our recent findings (<u>Karim et al., 2026</u>). The average [^18^F]MK-6240 GM/WM ratios were: DSAD= 2.82, AD= 2.45 and CN=1.81. The cases in the two studies were the same, but some variation in the brain sections was inevitable. The higher average ratio in CN subjects for both [^18^F]flortaucipir and [^18^F]MK-6240 was driven by an outlier. Figure-6C compares [^18^F]flortaucipir and [^18^F]MK-6240 in all the DSAD and AD subjects combined. Average ratio of [^18^F]MK-6240 was approximately 50% of the average ratio of [^18^F]flortaucipir.

**Fig. 6.**
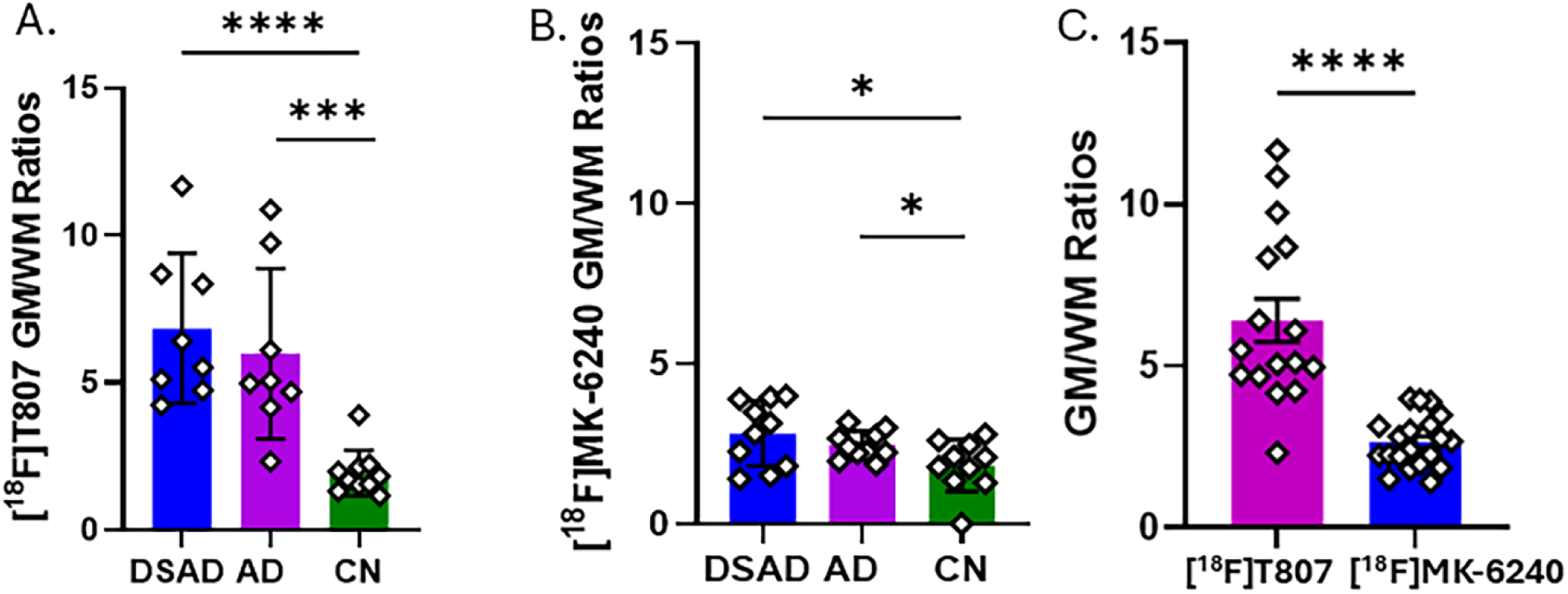
Group Comparisons of [^18^F]Flortaucipir and [^18^F]MK-6240 Binding: (A). [^18^F]Flortaucipir binding ratio, GM/WM of DSAD, AD, and CN cases. (B). [^18^F]MK-6240 binding ratio, GM/WM of DSAD, AD, and CN cases. (C). Comparison of [^18^F]Flortaucipir and [^18^F]MK-6240 GM/WM binding of all DSAD and AD cases. Unpaired two-tailed parametric t-tests determined statistical significance between each parameter (* p < 0.05, *** < 0.001, **** < 0.0001).

#### 3.2.6. Comparison of [^18^F]flortaucipir and [^18^F]FAZIN3 in PD

Binding of [^18^F]flortaucipir was compared in human post-mortem AC and corpus callosum (CC) from CN and PD cases previously evaluated using anti-ubiquitin immunohistochemistry and correlated with MAO-A imaging agent, [^18^F]FAZIN3 (<u>Mukherjee et al., 2022</u>). Figure-7 shows CN and PD cases (Fig-7C,G,K, O) in which cortical AC regions exhibited extensive amounts of [^18^F]FAZIN3 binding which was blocked by the selective MAO-A inhibitor, clorgyline (Fig-7Q). Neighboring sections from the same CN and PD cases did not show any significant [^18^F]flortaucipir binding (Fig-7Q). Ratio of [^18^F]FAZIN3 binding, total/ clorgyline=6 in the PD cases, while [^18^F]flortaucipir ratio was ∼1.3 (Fig-7Q).

**Fig. 7.**
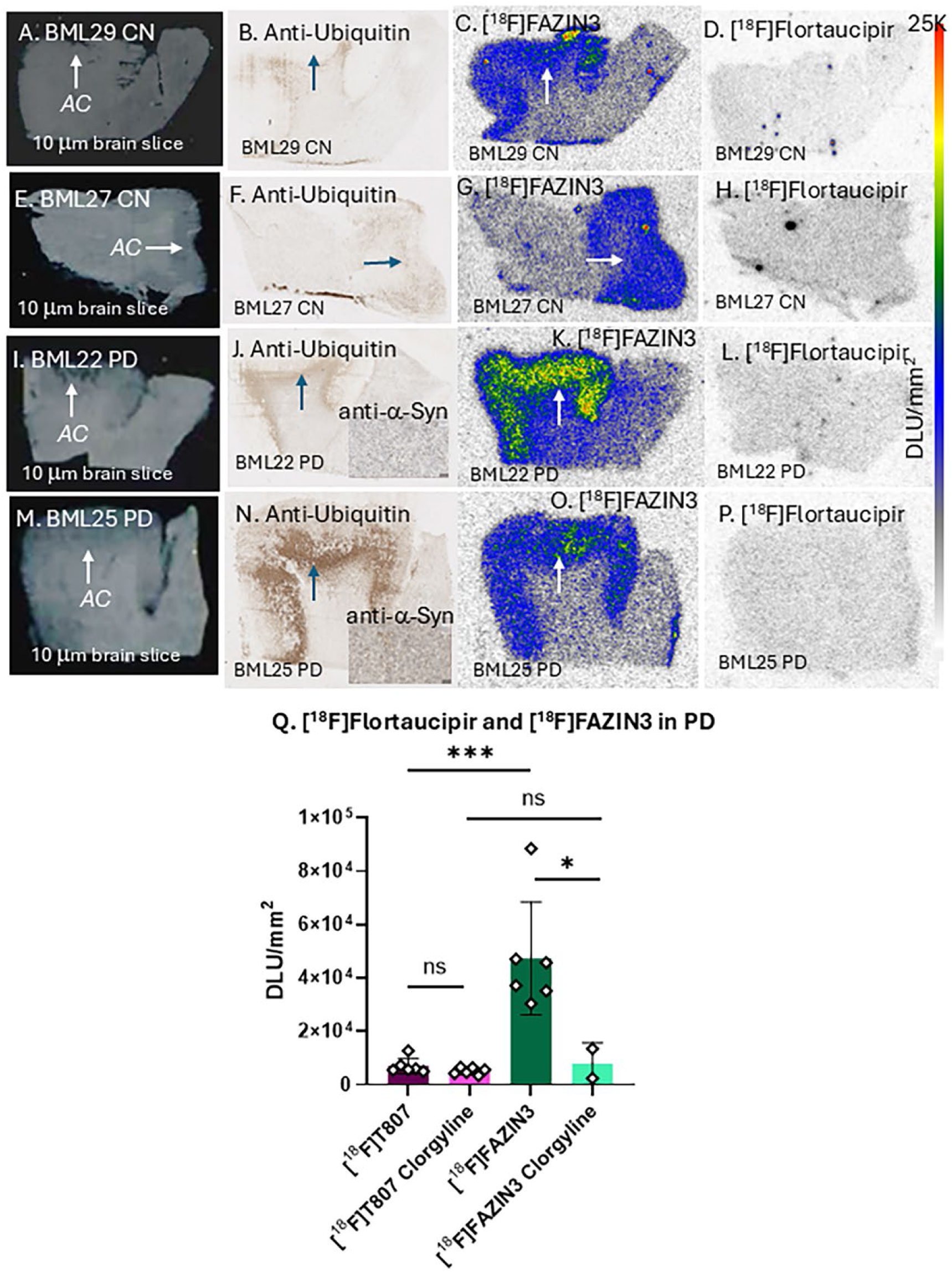
Comparison of [^18^F]FAZIN3 and [^18^F]Flortaucipir binding in CN and PD: (A, E, I, M): Postmortem human brain slice (10 μm) of CN (BML29 and BML27) and PD (BML22 and BML25) showing anterior cingulate (AC, shown by arrow); (B, F, J, N): Anti-ubiquitin for Lewy body, inset in J,N show anti-α-synuclein; (C, G, K, O): [^18^F]FAZIN3 cortical binding to MAO-A in AC of CN and PD sections; (D, H, L, P): Absence of [^18^F]flortaucipir cortical binding in AC of CN and PD sections; (Q). Plot shows binding of [^18^F]flortaucipir and [^18^F]FAZIN3 in PD cases without and with 10 μM clorgyline (* p < 0.05, *** < 0.001; ns: not significant).

## 4. Discussion

[^18^F]Flortaucipir (Tauvid^TM^) and [^18^F]MK-6240 (Tauklarify^TM^) are the two approved clinical tau-PET imaging agents for NFT evaluation in AD. Both imaging agents have been used in several PET studies, including clinical trials. A few head-to-head comparison studies have been reported during the last few years in order to gain a solid understanding of their tau binding characteristics in relation to the stage of the disease. As may be expected, due to interplay of the complex neuropathology in AD during the disease stages, pharmacology of the two radiotracers, in vivo binding selectivity, in vivo kinetics and other nonspecific binding issues, some differences in their tau detection capabilities have been reported. Aside from the off-target issues with both the radiotracers, the two main issues have been 1. Regional variation in tau binding in the brain, and 2. Earlier detection of tau in some brain regions.

**Table 2:** Tau binding affinities of Flortaucipir and MK-6240.

|  | **Flortaucipir** | **MK-6240** | **Ratio<sup>1</sup>** | **Radioligand** | **Reference** |
|---|---|---|---|---|---|
| Tau IC<sub>50</sub>, nM | 2.91±0.17 | 5.99±0.95 | 2.06 | [<sup>125</sup>I]IPPI | <sup>2</sup>Biju et al., 2026<br>This work |
|  | na | 4.5 (2.7 pM) | na | [<sup>3</sup>H]PI2620 | <sup>3</sup>Malarte et al., 2023 |
|  | 2 (0.1 pM) | 12 (1 pM) | 6 | [<sup>3</sup>H]MK-6240 | <sup>4</sup>Malarte et al., 2021 |
| Tau, Ki, nM | na | 0.36 | na | [<sup>3</sup>H]NFT-355 | <sup>5</sup>Hostetler et al., 2016 |
| Tau, Kd, nM | 14.6 | na | na | [<sup>18</sup>F]T807 | <sup>6</sup>Xia et al., 2013 |
|  | na | 0.32 | na | [<sup>3</sup>H]MK-6240 | <sup>4</sup>Malarte et al., 2021 |
|  | 1.71 | 0.26 | 0.15 | [<sup>3</sup>H]MK-6240<br>[<sup>3</sup>H]AV-1451 | <sup>5</sup>Hostetler et al., 2016 |
|  | 0.7 | na | na | [<sup>18</sup>F]AV-1451 | <sup>7</sup>Ono et al., 2017 |
|  | 0.4 | na | na | [<sup>3</sup>H]AV-1451 | <sup>8</sup>Vermeiren et al., 2018 |
|  | na | 0.42 | na | [<sup>3</sup>H]MK-6240 | <sup>9</sup>Walji et al., 2016 |
|  | 1.9 | 0.57 | 0.3 | [<sup>3</sup>H]MK-6240<br>[<sup>3</sup>H]AV-1451 | <sup>10</sup>Lindberg et al., 2021 |

<sup>1</sup>Ratio = MK-6240/Flortaucipir; <sup>2</sup>Affinity measured using AD brain slices with tau labeled with [<sup>125</sup>I]IPPI; <sup>3</sup>Affinity measured using AD brain homogenates with tau labeled with [<sup>3</sup>H]PI2620; <sup>4</sup>Affinity measured using AD brain temporal cortex homogenates with tau labeled with [<sup>3</sup>H]MK-6240 (in parenthesis, a super high affinity site was identified); <sup>5</sup>Affinity measured using AD brain homogenates with tau labeled with [<sup>3</sup>H]NFT-355/ [<sup>3</sup>H]MK-6240/ [<sup>3</sup>H]AV-1451; <sup>6</sup>Affinity measured using AD brain slices and autoradiography with tau labeled with [<sup>18</sup>F]T807; <sup>7,8</sup>Affinity measured using AD brain homogenate with tau labeled with [<sup>18</sup>F]T807 and affinity measured for MAO; <sup>6</sup>Affinity measured using AD brain slices and autoradiography with tau labeled with [<sup>18</sup>F]T807; 7 ; 8 ;9; 10 ; 11 ; na = not available; <sup>9</sup>Affinity measured using AD brain homogenates with tau labeled with [<sup>3</sup>H]MK-6240; <sup>11</sup>Affinity measured using AD brain homogenates with tau labeled with [<sup>3</sup>H]MK-6240/ [<sup>3</sup>H]AV-1451; na: not available.

There have been several reports on the measurement of binding affinity of the two radiotracers. Table-2 summarizes the finding of each of them with their reported values of either IC50, Ki (inhibition constant) or Kd (equilibrium dissociation constant). There have been 7 reports on the measurement of Kd and four of them show MK-6240 to have lower Kd values compared to flortaucipir. One report of Ki measures of MK-6240 shows it to have a sub-nanomolar affinity (<u>Hostetler et al., 2016</u>). Using [^3^H]PI260 for IC50 measurement, a value of 4.5 nM has been reported. In brain homogenates (<u>Malarte et al., 2023</u>). Another report using the same radioligand [^3^H]MK-6240 in brain homogenates for both MK-6240 and flortaucipir, flortaucipir was shown to have six-times higher affinity compared to MK-6240 (<u>Malarte et al., 2021</u>). Because of the limited head-to-head binding affinity measurement using the same radioligand under near-identical conditions, our results using [^125^I]IPPI binding in intact AD brain slices showed that flortaucipir has two times higher affinity compared to MK-6240. Molecular modeling studies on tau fibrils show similar binding capabilities of both flortaucipir and MK-6240 (<u>Mukherjee et al., 2021</u>; <u>Murugan et al., 2018</u>). Recent cryo-EM studies have revealed a 1:1 fibril to MK-6240 binding (<u>Kunach et al., 2024</u>). Similar binding of flortaucipir with tau fibril is expected based on molecular modeling studies.

Based on our recent report on the binding of [^18^F]MK-6240 in DSAD and AD subjects (<u>Karim et al.,</u> <u>2026</u>), we carried binding studies of [^18^F]flortaucipir using brain sections in the same cases using similar protocols. Extensive binding of [^18^F]flortaucipir was observed in the FCX and TCX of DSAD and AD cases. A distinct difference was the increased binding of [^18^F]flortaucipir compared to [^18^F]MK-6240 in the tau rich regions of the cortical areas. This resulted in a higher GM/WM ratio for [^18^F]flortaucipir in both the DSAD and AD cases. Average GM/WM ratio of [^18^F]MK-6240 was approximately 50% of the average ratio of [^18^F]flortaucipir. PET studies have reported a higher SUV for [^18^F]flortaucipir in AD subjects compared to [^18^F]MK-6240 (<u>Flores et al., 2020</u>; <u>Zammit et al., 2026</u>). The higher binding of [^18^F]flortaucipir may be related to the higher tau affinity of flortaucipir compared to MK-6240. Reported measured lipophilicity of flortaucipir was log P= 1.67 (<u>Xia et al., 2013</u>) and MK-6240 was logD= 3.32 (<u>Walji et al., 2016</u>). However, calculated lipophilicities, ClogP, using ChemDraw were flortaucipir= 3.18 and MK-6240= 2.76, suggesting flortaucipir to be more lipophilic compared to MK-6240. The difference between the lipophilicity of the two radiotracers may potentially play a role in SUVR measures. Differences in binding affinity and lipophilicity amongst radiotracers can translate to higher target-to-nontarget ratios which has been observed in various neuroreceptor PET studies (<u>Mukherjee et al., 1995</u>; <u>Mukherjee et al., 2002</u>).

Since [^18^F]flortaucipir has been reported to have an affinity for MAO (higher affinity for MAO-A compared to MAO-B), there has been speculation if part of the difference between [^18^F]flortaucipir and [^18^F]MK-6240 may be related to this additional non-tau binding. However, as mentioned earlier, in vivo MAO-A or MAO-B binding has not been established for [^18^F]flortaucipir. Using postmortem brain sections of CN and PD subjects, we show absence of MAO-A binding in the anterior cingulate. Brain sections from these subjects were previously shown to bind extensively to the selective MAO-A radiotracer, [^18^F]FAZIN3 (Mukherjee et al., 2022). Experiments with [^18^F]flortaucipir require alcohol washing conditions, while [^18^F]FAZIN3 binding and washing protocols used only PBS. Previous studies to ascertain MAO-A binding of [^18^F]flortaucipir using alcoholic and nonalcoholic in AD brain sections were unsuccessful in confirming MAO-A binding (<u>Drake et al., 2019</u>; <u>Hansen et al., 2018</u>; <u>Wright et al., 2022</u>). We have previously reported an increase in MAO-A using [^18^F]FAZIN3 in the anterior cingulate and its relationship with Aβ plaques and tau in the same cases (Syed et al., 2023). An interplay between MAO-A/Aβ+/tau+ cannot be ruled out. It remains unclear whether this MAO-A/Aβ+/tau+ complex, if formed, will be able to bind to [^18^F]flortaucipir, while [^18^F]MK-6240, which does not have an affinity for MAO-A will not bind.

More recent studies also indicate the ability of [^18^F]MK-6240 to detect low levels of tau while [^18^F]flortaucipir is unable to (<u>Gogola et al., 2022</u>; <u>Povola et al., 2026</u>). This is paradoxical based on binding affinities and reported lipophilicities. It is suggested that differences in Kd between [^18^F]MK-6240 and [^18^F]flortaucipir may account for this. Table-1 shows equilibrium dissociation constants, Kd for flortaucipir in the range of 0.4 to 1.9 nM, with one outlier of14.6 nM, while MK-6240 ranges from 0.26 to 0.57 nM. Thus, based on Kd measurements, MK-6240 has 3 times higher affinity compared to flortaucipir. However, the lower IC50 measurements for flortaucipir compared to MK-6240 favor higher binding of [^18^F]flortaucipir. Molecular modeling shows 4 high binding sites for flortaucipir and MK-6240 with similar binding energies, with flortaucipir being more favorable (<u>Mukherjee et al., 2021</u>; <u>Murugan et al., 2018</u>). CryoEM of fibril tau with bound MK-6240, exhibited a 1:1 binding at tau Site 1 (<u>Kunach et al., 2024</u>). Since PET studies of [^18^F]flortaucipir and [^18^F]MK-6240 are carried out with high specific activity (low mass), it is very likely that only Site 1 is being occupied by the two radiotracers in PET studies (<u>Kunach et al., 2024</u>). The higher GM binding of [^18^F]flortaucipir compared to [^18^F]MK-6240 in this work and the higher SUV of [^18^F]flortaucipir in PET studies, all point to the higher affinity of flortaucipir to tau. In the case of both flortaucipir and MK-6240, a very high affinity binding site with picomolar affinity has been suggested based on tissue homogenate binding assays (Table-2, <u>Malarte et al., 2021</u>; <u>Malarte et al., 2023</u>). Aside from binding affinity to tau, SUVR in PET studies may be influenced by the chosen reference region and indirectly on the lipophilicity of the radiotracer. From our calculated log P, it appears that flortaucipir is more lipophilic compared to MK-6240. Thus, the higher SUVR of [^18^F]MK-6240 and its ability to detect tau earlier may be due to the differences in nonspecific binding in the reference non-tau regions of [^18^F]MK-6240 (<u>Pascoal et al., 2018</u>) compared to [^18^F]flortaucipir (<u>Baker et al., 2017</u>; <u>Barret et al., 2017</u>; <u>Wooten et al., 2017</u>).

Limitations of the study include the small number of DSAD and AD cases in advanced Braak stage, with minimal diversity in tau abundance and disease severity. Cases were not age matched between different groups, so CN cases were the oldest while DSAD and AD cases were more similar in age. Variability across brain sections in the three groups for close comparisons of radiotracer binding.

In summary, our results suggest that binding affinity of flortaucipir and MK-6240 are within the same nanomolar range for AD tau in brain slices, with flortaucipir being slightly more potent than MK-6240. Because these values were measured in intact brain slices, they are likely reflective of tau affinity in the whole brain. Higher GM/WM ratios in DSAD and AD cases of [^18^F]flortaucipir compared to [^18^F]MK-6240 are consistent with higher SUV for [^18^F]flortaucipir in PET studies. This study was unable to detect any off-target binding of [^18^F]flortaucipir in the anterior cingulate of CN and PD cases, particularly to MAO-A.

## Author Contributions

All authors had full access to evaluate the integrity of the data and the accuracy of the data analysis. Study concept and design: JM. Acquisition of data: APB, JMH, CCM, EMH, CL, and JM. Analysis and interpretation of data: APB, JMH, CCM and JM. Drafting of the manuscript: APB, JMH, EMH and JM. Statistical analysis: APB, JMH, CCM. Obtained funding: JM. Study supervision: JM.

## Institutional Review Board Statement

Not applicable

## Informed Consent Statement

Not applicable.

## Data Availability Statement

The data that support the findings of this study are available from the corresponding author upon reasonable request.

## Acknowledgments

Research support provided by NIH AG R01 AG077700 (JM) and the Undergraduate Research Opportunities Program (UROP) at University of California, Irvine. We are grateful to Dr. Elizabeth Head at the UCI MIND Institute for brain tissue used in the current study which was supported by funding from the UCI ADRC grant NIH/NIA P30AG066519. The UCI ADRC is grateful to the donors and families who participated in the gift of brain donation. We are also grateful to Dr. Geidy Serrano of the Banner Sun Health Research Institute Brain and Body Donation Program of Sun City, Arizona for the provision of brain tissue. The Brain and Body Donation Program is supported by the National Institute of Neurological Disorders and Stroke (U24 NS072026, National Brain and Tissue Resource for Parkinson’s disease and related disorders), the National Institute of Aging (P30 AG19610 and P30AG072980, Arizona Alzheimer’s Disease Center), the Arizona Department of Health Services (contract 05700, Arizona Alzheimer’s Research Center), the Arizona Biomedical Research Commission (contracts 4001, 0011, 05-901 and 1001 to the Arizona Parkinson’s Disease Consortium) and the Michael J. Fox Foundation for Parkinson’s Research. We thank Jeffrey Kim and Delia Tifrea in Pathology and Laboratory Medicine, University of California-Irvine for immunostaining of brain sections. We thank Lena Qin for brain tissue sectioning and Fariha Karim for technical assistance. We acknowledge the use of QuPath for digital analysis of IHC.

